# Differential transport of a guild of mutualistic root aphids by the ant *Lasius flavus*

**DOI:** 10.1101/2022.07.05.498828

**Authors:** Thomas Parmentier

## Abstract

Mutually beneficial associations are widespread in ecological networks. They are typically assembled as multispecies guilds of symbionts that compete for one or more host species. The ant *Lasius flavus* engages in an intriguing and obligate mutualistic association with a community of aphids that are cultivated on plant roots in its nests. The ant displays a repertoire of amicable behaviours towards the aphids, including their transport. I examined whether *L. flavus* preferentially carried some of the root aphids. Using a no-choice and a choice experiment, I comparatively analysed the transport rate of five obligate and one loosely associated species back to the ant nest and used the transport rate of the ant larvae as a reference. All associated root aphids were carried back to the nest, but in a clear preferential hierarchy. *Geoica utricularia, Forda Formicaria* and *Trama rara* were rapidly transported, but slower than the own larvae. *Tetraneura ulmi* and *Geoica setulosa* were collected at a moderate rate and the loosely associated *Aploneura lentisci* was slowly retrieved. In contrast, different species of unassociated aphids were not transported and even provoked aggressive behaviour in *L. flavus*. This study revealed that co-occurring symbionts may induce different degrees of host attraction, which ultimately may affect the coexistence and assembly of ant-symbiont communities.

## 1. INTRODUCTION

Reciprocally beneficial or mutualistic interactions have been traditionally studied as one-to-one relationships between two partner species. However, multiple symbionts often compete for the beneficial services of one or more partner species at the same time (Stanton 2003; Palmer et al. 2012). Recent research gradually tries to grasp the complexity of the interactions within such guilds of co-occurring symbionts. These studies hinted that, in line with the well-known coexistence mechanisms within trophic guilds, competitive coexistence of mutualist guilds may be facilitated by processes such as competition-colonization trade-offs (Yu et al. 2004), niche differences (Sampayo et al. 2007; Peay 2016) and indirect interactions (Lee and Inouye 2010; Martignoni et al. 2020).

Ants have been an exquisite model group to study the ecology of symbiotic networks (Ivens et al. 2016). They are dominant arthropods that engage in an unparalleled diversity of symbiotic associations (Hölldobler and Wilson 1990; Parmentier 2020). Ant workers carry a whole range of items in, to and away from the nest including prey, seeds, leaves, nest material, and live and dead nest mates. The brood is also carried around in the nest or evacuated after disturbance (Hölldobler and Wilson 1990). Interestingly, some parasitic associates, including beetles and the *Phengaris* caterpillars, can also be picked up and brought as Trojan horses to the brood chambers or food storages (Hölldobler 1967; Cammaerts 1999; Solazzo et al. 2012; Parmentier 2019, 2020). Not only parasites, but also mutualistic aphids are picked up by some ants (Donisthorpe 1927; Way 1963; Heie 1980). As such, these symbionts can rapidly be moved to food plants or brought to safety.

The yellow meadow ant *Lasius flavus* (Fabricius, 1782) is a widespread Palearctic ant that lives in underground nests in grassland habitats (Seifert 2007; AntWiki 2022). *Lasius flavus* colonies are completely dependent on root aphids which are kept in high numbers in nest chambers built around herbaceous and grass roots. The root aphids gregariously feed on the sap in the roots and secrete droplets of sugary honeydew. This honeydew appears to be the main food source of a *L. flavus* colony. Different species of obligatory ant-associated root aphids co-occur in a *L. flavus* nest (Nielsen et al. 1976; Pontin 1978; Godske 1992; Depa and Wegierek 2011). These obligatory ant-associated root aphids evolved to a life in strict association with their ant host which was accompanied with behavioural (e.g., retracting appendages before transport, Bilska et al. 2018) and morphological adaptations (Depa et al. 2020). The association between root aphids and *L. flavus* is extremely intimate. The root aphids are licked, cleaned, and are also carried around by the *L. flavus* host when the nest is opened (Donisthorpe 1927; Way 1963; Paul 1977; Fig. 1). Root aphids can also be found in nests of other ants (Parmentier et al. 2020). These ants typically have a lower dependency on the root aphids (AntWiki 2022).

**Fig. 1.**
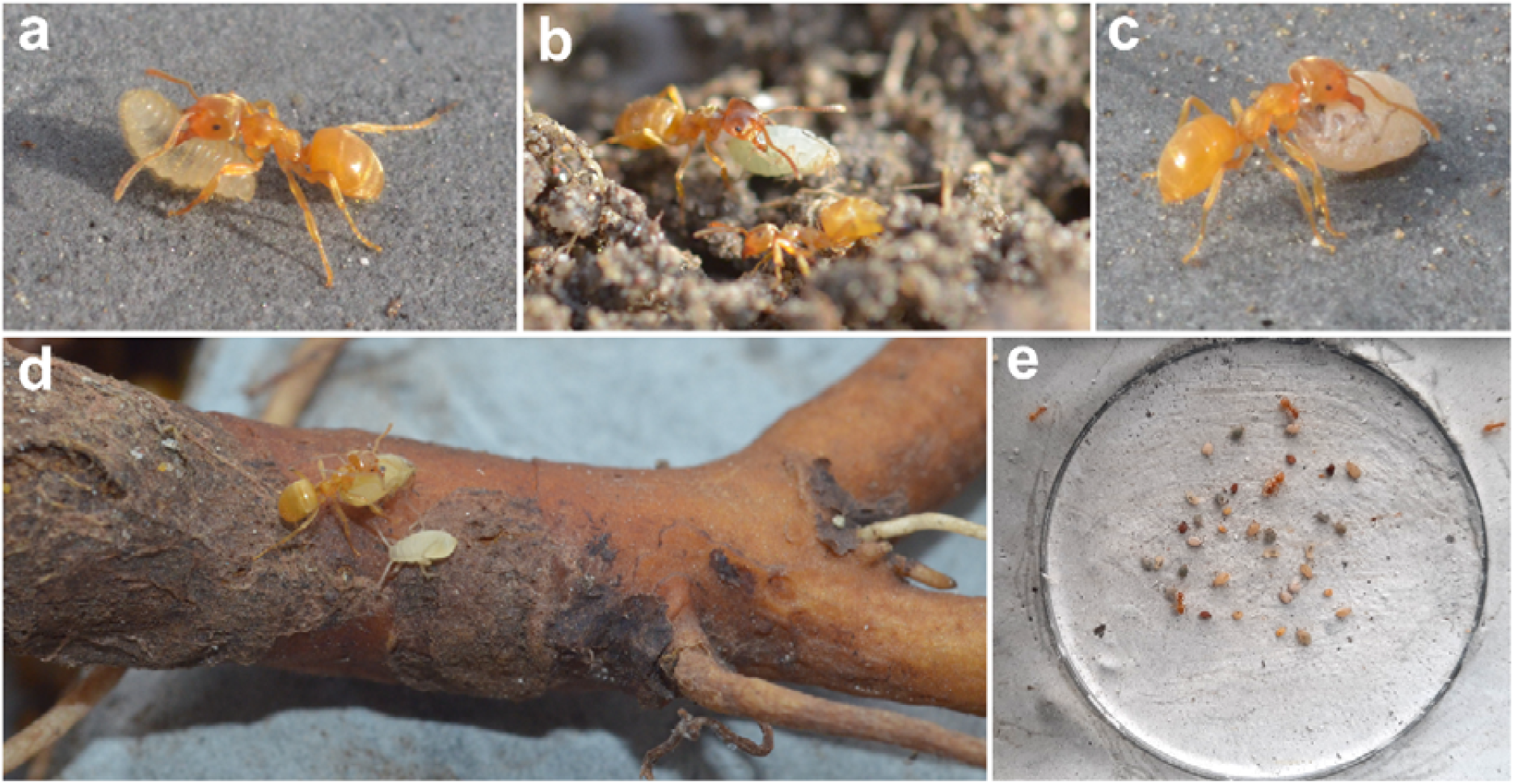
Overview of the interaction between *Lasius flavus* and its associated root aphid guild. A *L. flavus* worker carrying a larva (a), workers transporting different root aphid species: *Forda formicaria* (b), *Geoica utricularia* (c) and *Trama rara* (d). An unattended nymph of *T. rara* is feeding on the *Taraxum officinale* root (d). Start of an experimental trial in the choice experiment with five larvae, and five individuals of each of the five root aphid species in the central arena (e).

Our current understanding of the behavioural interactions between ants and root aphids is mostly descriptive (Zwölfer 1958a, b; Way 1963; Heie 1980), only a few studies quantified the behavioral interaction between ants and root aphids (Salazar et al. 2015). A first attempt to analyse the behaviour of *L. flavus* towards different root aphids more quantitatively was done in (Ivens 2012), albeit this study was preliminary in nature (Box A in (Ivens 2012)). This study found no major differences in ant behavioural repertoire towards different root aphids. Moreover, the aphids in the behavioural assays were poorly attractive which is likely caused by the minimalistic test arenas with only two workers.

Several studies already compared the ant visitation preference for different co-occurring aboveground aphids (Fischer et al. 2001; Woodring et al. 2004; Akyildirim et al. 2014; Pålsson et al. 2020). Highly visited aphids are typically better protected against enemies (Fischer et al. 2001). Carried to safety or to food plants (Way 1963) by the host may confer much higher benefits for the aphids than mere visitation and enemy deterrence, but no studies have comparatively examined this strong, mutualistic behaviour yet. Here, I compared the rate of transport to the nest of six root aphid species (five obligatory myrmecophilous, one loosely associated) by the host ant *L. flavus*. In addition, I compared their retrieval rate with those of the host’s own larvae, enabling us to assess whether they prefer some aphid partners over their own kin. Lastly, I offered five unassociated aphid species to test whether the ants showed amicable behaviour and provided transport services to unfamiliar aphid species.

## MATERIALS AND METHODS

### Study species

I focused on six species of root aphids living in the nest of *L. flavus*. Five species of the community of associated root aphids are considered as obligatory ant-dependent (Heie, 1980; Way, 1963), i.e. *Geoica setulosa* (Passerini, 1860), *Geoica utricularia* (Passerini, 1856), *Forda formicaria* von Heyden, 1837, *Trama rara* Mordvilko, 1908 and *Tetraneura ulmi* (Linnaeus, 1758). The obligate ant association of these species is also echoed in different morphological traits. The sixth species, *Aploneura lentisci* (Passerini, 1856), is characterized by a loose association with ants (Paul 1977). It can be found in ant nests, but mostly in the soil away from ant nests (Donisthorpe 1927; Paul 1977). I also collected five aphid species that are not associated with *L. flavus* at different sites in Northern Belgium: *Aphis sambuci* Linnaeus, 1758, *Cavariella aegopodii* (Scopoli, 1763), *Cinara laricis* (Hartig, 1839), *Macrosiphum rosae* (Linneaus, 1758) and *Periphyllus testudinaceus* (Fernie, 1852). As these aphids live on aboveground plant parts, and *L. flavus* does not forage aboveground, these species normally do not interact. However, *A. sambuci, C. laricis* and *P. testudinaceus* are frequently visited by aboveground foraging ants. During sampling the first two species were heavily visited by *Lasius niger* and the latter by *Formica polyctena. Cavariella aegopodii* and *M. rosae* are only occassionally tended by ants (Dhatwalia and Gautam 2009; Akyürek et al. 2016) and were not visited by ants at the time of sampling. Aphid identification was checked by different keys and guides (Heie 1980; Blackman and Eastop 2018; Blackman et al. 2019). Species id of the used aphids was individually verified after experimental trials.

### Aphid transport in a no-choice experiment

With this experiment, I compared the transport rate of co-occurring root aphids by offering single individuals to their ant host colony. I checked how many interactions were needed to trigger carrying behavior. For this experiment, I sampled ten different *L. flavus* nests in urbanized grassland sites near Ostend, Belgium (Fig. S1) (March-April 2022). Nests were selected that housed at least three species of the six focal root aphids (overview of the collected species per nest see Table 1). When the nest was opened, I frequently observed ants transporting different root aphids into safety (Fig. 1). In contrast to Ivens et al. (2012) that found a single aphid species in more than 50% of the sampled *L. flavus* nest, the tested root aphid species typically occurred together in the nest (cohabiting species, see also Godske 1992; Depa and Wegierek 2011). *Trama rara* was found on the roots of *Taraxum officinale*, the other aphids on grass roots. Root aphids were carefully taken from the roots in the nest and stored in a plastic container with plaster bottom. Workers of the host colony, some roots and nest material were also added. From each ant nest, 1000 workers and 150 larvae were separated and housed in a plastic box (27×8.4×9 cm) with a plaster bottom and fluon coated walls. On both ends of the box, nest sites were made, which were circular cavities (diameter 55 mm, depth 10 mm) in the plaster covered with a piece of cardboard. The ants readily brought their larvae in these nest sites and gathered around. Two separate nest sites in the box were chosen to promote ant traffic in-between. Central in the box, there was a circular arena which was made from a plaster filled petri dish (diameter 55 mm). The top rim of this dish was even with the plaster bottom of the box. The dish was filled with plaster to ca. 1 mm from the top rim resulting in a small plastic border surrounding the dish. After one day acclimatization of the ants to the lab box, an experimental trial was started by placing a root aphid individual that was collected in the same colony in the central arena. Then I counted the number of interactions with workers needed to initiate carrying behavior to the covered nest sites. An aphid-ant interaction took place when the ant antenna touched or crossed the body of the aphid. Although some workers could have engaged in more than one interaction if they returned to the aphid, consecutive interactions generally took place with unique workers. A trial was stopped when the root aphid was transported. The transporting ant and aphid were removed before they reached the nest site. If the aphid was not transported after ten interactions, it was removed, and the trial was also stopped. This methodology was followed in the subsequent trials with different root aphid individuals (aim was to have around ten unique individuals of each species, details see Table 1). These individuals belonged to different root aphid species, that were collected in the same nest as the ant workers of the lab nests. The order of the tested root aphid individuals was randomized. Root aphids were not re-used in subsequent trials. There was a pause of one minute between different trials. Interactions in subsequent trials were typically with a different set of workers, as transporting workers of previous trials had been removed and because of a constant and steady flow of workers going from one nest site to the other. As a reference, I also checked the number of interactions needed to transport the own larvae using the same setup. Trials were conducted under ambient light and at room temperature (20±1°C).

**Table 1.**
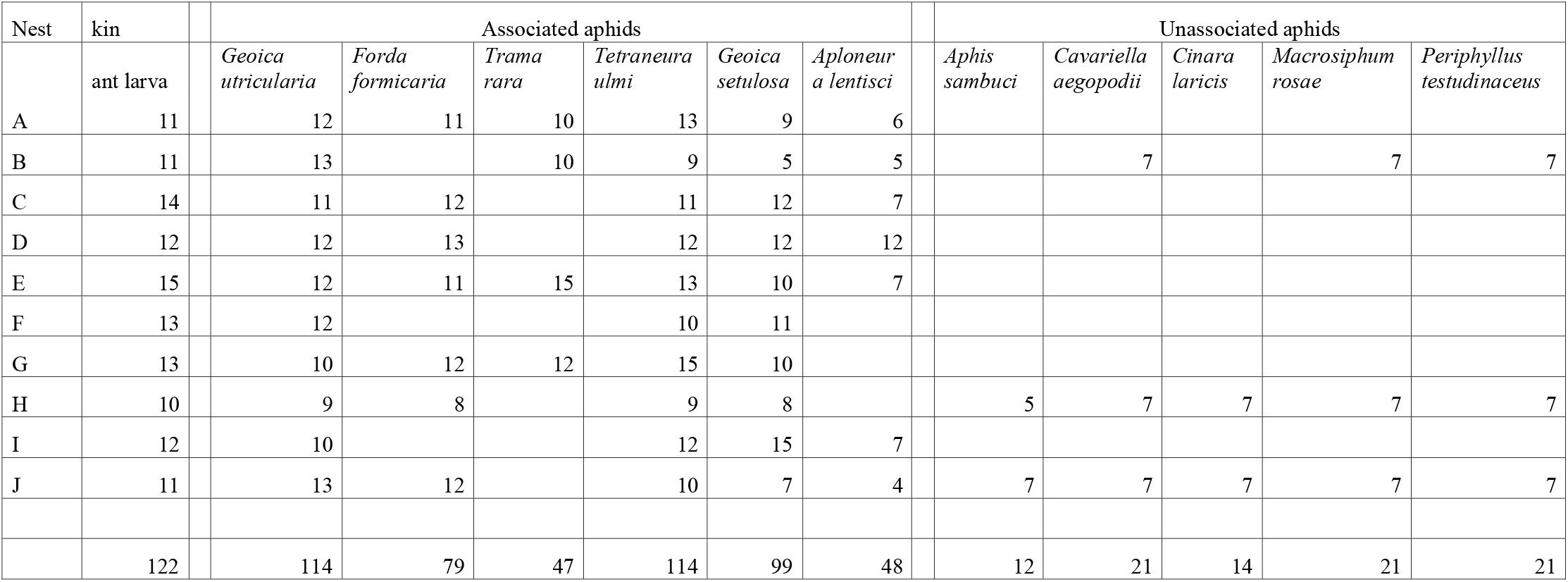
Number of unique individuals (ant larva or aphids) tested in the no-choice experiment for the ten experimental *L. flavus* colonies. The location of the ten *L. flavus* nests (A-J) are given in Fig. S1. The associated aphids and host workers used in a trial were collected in the same *L. flavus* nest.

I compared the retrieval rate of the six associated aphids and larvae with a mixed-effects Cox proportional hazards model using the coxme function in the coxme package (Therneau 2015). This type of model models time-to-event data (in this case number of interactions until carrying) and allows the inclusion of right-censored data, which are in this experiment aphids or larvae that were not collected and remained in the central arena after ten interactions. Species (7 levels: 6 aphid species and the host larvae) was modeled as fixed factor, host colony (10 levels) as a random factor. The proportional hazards assumption was met (proportional hazards test with the cox.zph function in package survival, Therneau 2020: species: χ2 = 4.53, df = 6, *P =* 0.61), The cumulative distribution of aphids or larvae retrieved over time (opposite of a survival curve) were plotted with the ggsurv function and were based on a proportional hazards models without the random factor colony (Therneau 2020).

In addition, I tested whether five unassociated aphid species triggered transport behavior using the same protocol, but only in two or three of the ant colonies (Table 1).

### Aphid transport in a choice experiment

In this experiment, the six focal root aphid species and the host larvae were presented to *L. flavus* colonies at the same time. The aim was to check whether some species were preferentially carried to the nest when alternative choices were possible. I dug out six independent colonies of *L. flavus* in a park in Ostend, Belgium (March-April 2021) (Fig. S1, minimum inter-nest distance 173 m). The six root aphids were collected from clusters of *L. flavus* nests at one neighbouring site (polygon in Fig. S1, area = 2180 m^2^). For each trial, I randomly scattered five individuals of each of the six root aphid species (= 30 root aphid individuals) and five host larvae in the central arena. Ants rapidly discovered the aphids and larvae and started to carry them back to one of the two nest sites. Ants could easily walk in and out the central arena with their load. Wandering root aphids were hindered to leave the central arena by the small plastic border. In case they could escape, they were gently placed back with a fine brush. The central arena was photographed at a two-minute interval over a total period of 30 min. Afterwards, I analyzed the photo sequence and counted the number of remaining individuals out of 5 for each of the six root aphid species and host larvae at each time point. Sometimes, aphids or larvae were picked up but dropped before leaving the arena. These individuals were not considered as retrieved. Afterwards I lifted the cardboard covering the nest sites and gently removed the root aphids. I also removed the aphids and larvae left in the central arena. I repeated this experiment 10 times for each colony (60 trials in total), for each trial a unique set of aphids was used. There was a period of 30 min between successive trials.

These data were also analyzed with a mixed-effects Cox proportional hazards model. Rather than the number of interactions, here the time (minutes) until carrying was of interest. The ants typically antennated different aphids or larvae before picking up an individual (video S1). In addition to species, the order of the choice trial in a test colony was incorporated as a continuous factor (1 = first trial to 10 = last trial in a colony), to assess whether the retrieval rate in a colony remained constant over the consecutive trials. The interaction between species and the order of the trial was also included. Colony (6 levels) was modelled as a random factor, trial (10 levels for each colony) was nested within colony. The constant hazard assumption was violated when a time interval of two minutes was taken. However, by surveying the transport of individuals every six minutes over the 30 minute time interval, model assumptions were met (cox.zph function: species: χ2 = 6.57, df = 6 *P =* 0.36, sequence: χ2 = 1.66, df = 1, *P* = 0.20, full model: χ2 = 8.16, df = 7, *P =* 0.32)

### Behavior of *L. flavus* towards aphid species

During the no-choice experimental trials with associated and unassociated aphids outlined above, I also categorized the different ant interactions: Apart from carrying behavior, I discriminated ignoring (= an ant encounters an aphid, but continues without any change in behavior), inspecting (= an ant detects an aphid, stops or turns its head to the aphid, but then moves on), antennating (= rapid drumming of the antennae to scan the aphid), opening of the mandibles (= threat posture, an ant opens its mandibles, but does not attempt to bite), biting (= an ant snaps with its mandibles), abdomen bending (= an ant bend its abdomen to spray formic acid, this behavior is accompanied by biting). Note that carrying was often preceded by heavily antennating, but this interaction was then categorized as carrying. The number of these non-carrying behaviors per trial varied from 0 (when the aphid was carried in the first interaction) to 10 (when no carrying occurred). Note that trials where the aphid was already carried in the first interaction, were not included as none of the focal behaviors then took place. The frequency of each type of non-carrying behavior in an ant-aphid interaction was compared among the ten aphid species (both associated and unassociated) with a Permanova (function “adonis,” 999 permutations, strata: colony). Next, I compared the proportion of antennating out of all non-carrying behaviors in the different aphid species. Here, I used a binomial generalized linear mixed model with the proportion of antennation as dependent variable and aphid species as fixed factor (package lme4, (Bates et al. 2016)) Host colony was added as a random factor. An observation level random factor was also modelled to account for overdispersion (Browne et al. 2005).

## RESULTS

### Aphid transport in a no-choice experiment

The yellow meadow ant carried the associated root aphids in a clear hierarchy in the no-choice experiment (Cox mixed-effects model: LR test: χ2 = 189.2, df = 6, *P* **<** 0.0001). *Trama rara, Geoica utricularia* and *Forda formicaria* were the three most preferred root aphid species (significances of Post hoc Tukey tests indicated with letter codes on Fig. 2a). More than half of the individuals of these species were already retrieved after three interactions (Fig. 2a). Most of the root aphids *Geoica setulosa* and *Tetraneura ulmi* were also carried back to the nest after 10 interactions, but the retrieval rate was slower than the three preferred species (half of individuals transported after six interactions). The loosely ant-associated root aphid *A. lentisci* was characterized by the lowest attraction and was often not carried after 10 interactions. The hazard ratios give the transport/hazard rate of the aphids relative to the transport/hazard rate of the ant larvae (Fig. 2a). Although the very high retrieval rate of the root aphids, host larvae were significantly more attractive (half of larvae carried after two interactions). Unassociated aphids were not picked up and transported to the nest.

**Fig. 2.**
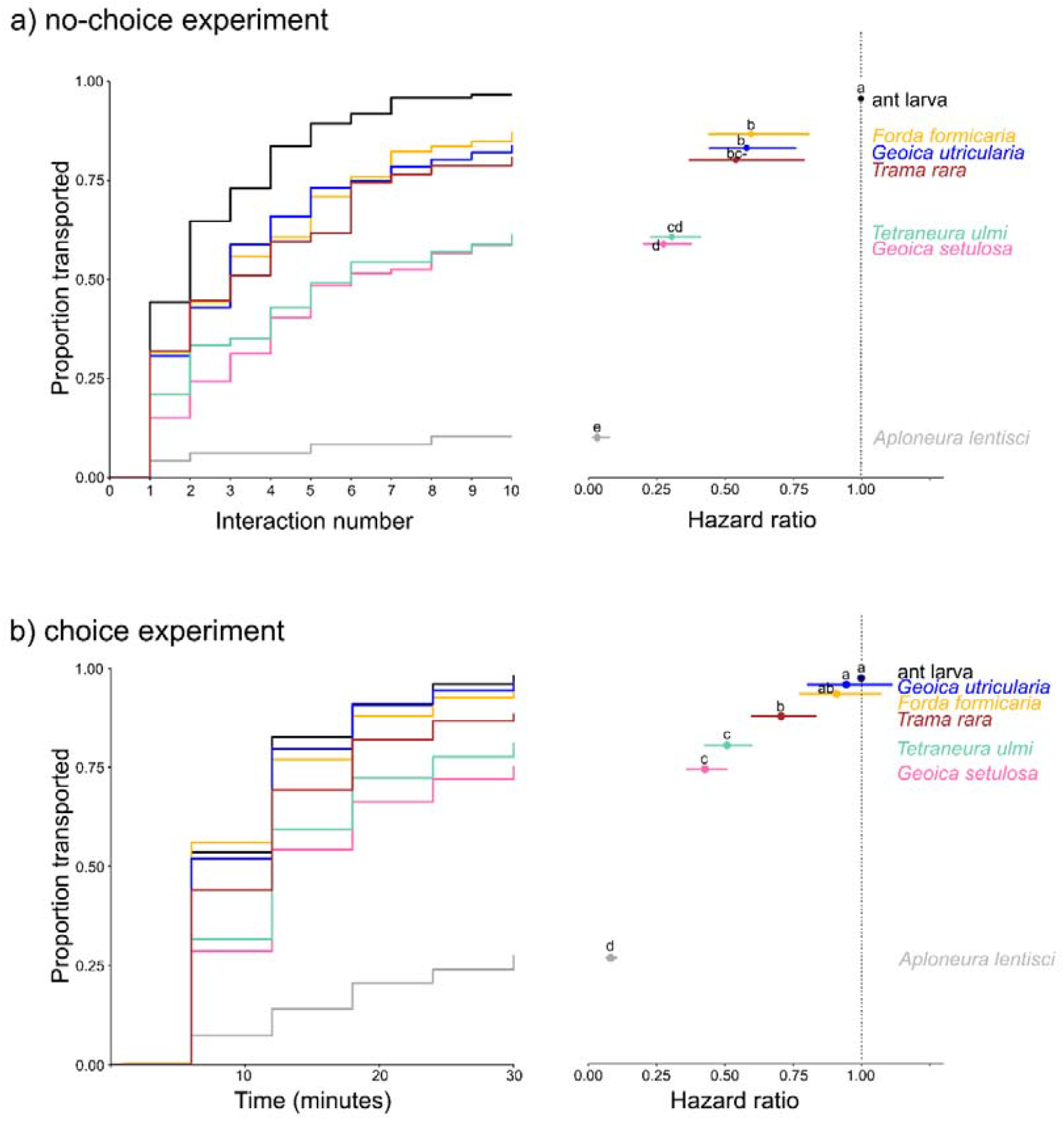
Preferential transport hierarchy of root aphids associated with *L. flavus*: a) no-choice experiment: trials with one aphid individual at the same time. Transport of the aphids followed over ten interactions with the host ant displayed in curve with the cumulative number of transport events. Corresponding hazard ratios (relative to the transport rate of the host larvae) are given on the right. b) choice experiment: six aphid species and larvae offered at the same time; transport determined over a time frame of 30 minutes (interval 6 minutes) displayed in curve with the cumulative number of transport events. Corresponding hazard ratios (transport rate relative to the transport rate of the host larvae) are given on the right. Hazard ratio = 1, transport rate similar to transport rate of own larvae, HR: 0.5 transport rate of aphid is half of the transport rate of the own larvae. Post-hoc differences in retrieval rate are indicated with a letter code. Hazard ratios that do not share a letter are statistically different (Post-hoc Tukey’s test, 5 % probability level).

### Aphid transport in a choice experiment

When offering the six associated root aphid species at the same time, a similar preferential hierarchy was found as in the no-choice experiment (Cox mixed-effects model with observations every six minutes: LR test: χ2 = 857.3, df = 6, *P* < 0.0001). *G*e*oica utricularia, F. formicaria* and *T. rara* were rapidly retrieved. After six minutes half of the individuals of *G. utricularia* and *F. formicaria* were transported, the median retrieval time for *T. rara* was eight minutes (Fig. 2b). *Geoica setulosa* and *Tetraneura ulmi* were transported at a modest rate (half of the individuals collected after 12 min., Fig. 2b), and *A. lentisci* was slowly retrieved and often not transported. The retrieved individuals of this species were also frequently dropped outside the nest, which was not observed in the other aphid species. Contrary to the no-choice experiment, I found no difference in the retrieval rate of the ant larvae (half collected after six min., Fig. 2b) and the three most preferred root aphids (significances of Post hoc Tukey tests on the hazard ratios indicated with letter codes on Fig. 2b). The proportion of individuals transported in the choice experiment declined in successive colony trials (Cox mixed-effects model: LR test: χ2 = 8.7, df = 1, *P* = 0.003).

### Behavior of *L. flavus* towards aphid species

The behavioral repertoire towards the six associated species was very amicable. If they were not transported, they were often antennated. They never provoked an aggressive response (opening mandibles, biting, spraying formic acid). Ants showed different levels of amicable behavior towards the guild of associated aphid species (PERMANOVA, df = 6, F = 20.8, *P* = 0.001) In general, species that were more rapidly transported, were also more antennated (Tukey Post hoc differences in proportion antennation indicated with letter code on Fig. 3). Ant behavior towards unassociated species was markedly different (PERMANOVA, df = 10, F = 20.6, *P* = 0.001). They showed aggressive behavior towards the unassociated species. They tried to bite them and in some cases they were dragged around. The level of provoked aggression depended on the aphid species, but it was striking that even obligatorily ant-associated aphids were strongly attacked.

**Fig. 3.**
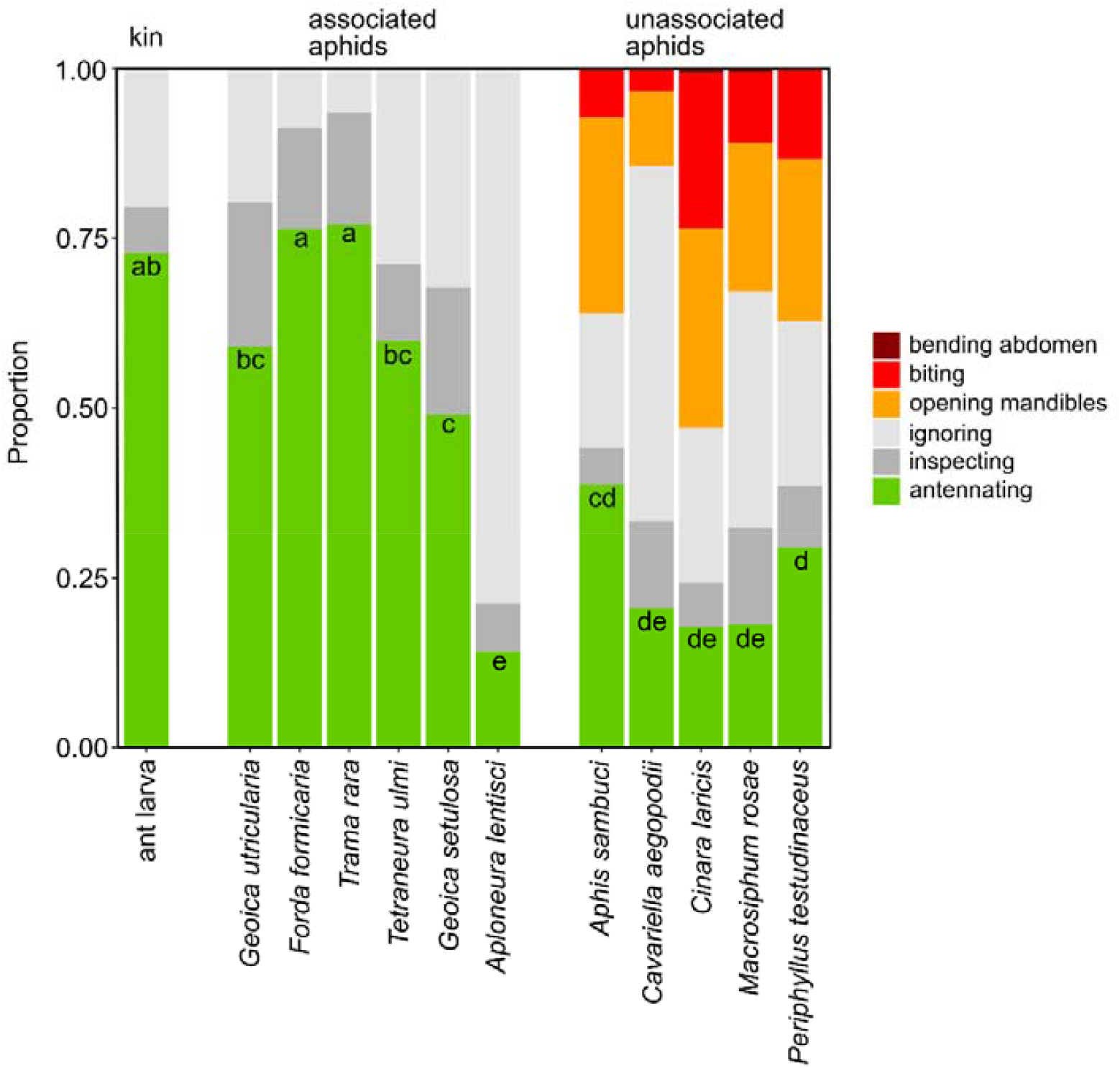
Behavioral repertoire of *L. flavus* workers interacting with associated root aphids and unassociated aphids. A letter code on top of the anntenation bars indicates whether species significantly differ in the proportion of antennation elicited (Post-hoc Tukey test, binomial GLMM)

## DISCUSSION

This study underscores the intimate association between the ant *Lasius flavus* and a community of associated root aphids (Zwölfer 1958b, a; Seifert 2007; Ivens 2012). Associated aphids were heavily antennated, rapidly picked up and brought back to the nest. However, the ant *L. flavus* showed different degrees of amicability towards the group of associated mutualists, which was demonstrated with a clear hierarchy in the transport rate of the aphids. The root aphids *Trama rara, Forda formicaria* and *Geoica utricularia* were faster retrieved than the two other obligate ant-associated root aphids *Tetraneura ulmi* and *Geoica setulosa*. As expected, the facultatively associated root aphid *A. lentisci* was the least attractive. Ants often ignored this aphid and carrying was infrequent.

The carrying behavior of *L. flavus* was clearly selective as they did not move any material that they encountered. Unassociated aphids were never transported and, like other ants, *L. flavus* showed aggression towards non-partner aphids (Sakata 1994; Hayashi et al. 2015). Threat postures and biting were never observed in the interaction with the six species of associated root aphids, even not in the interaction with the loosely associated *A. lentisci*.

Intriguingly, the three most preferred aphids elicited an equal attraction response than the own kin (larvae) in a choice experiment. This underlines the very high value of these aphids for the colony functioning (Stadler and Dixon 2005; Ivens 2012; Ivens et al. 2018). Nevertheless, when there was no choice, host larvae were carried faster than the most preferred root aphids.

Preference hierarchies have already been demonstrated in many other mutualisms, including pollination and plant mycorrhiza symbioses (Sanders 2003; Phillips et al. 2020). Gradations in partner attractivity has been tested in ant mutualisms as well. Workers of an ant colony show marked preferences for seeds of myrmecochorous plants (Leal et al. 2014; Miller et al. 2020) or for tending lycaenid caterpillars (Stadler et al. 2003). Ant visitation of aboveground aphids is also not random, with some aphids being much more tended than others (Fischer et al. 2001; Woodring et al. 2004; Akyildirim et al. 2014; Pålsson et al. 2020). Interestingly, the quality of honeydew secreted by aboveground aphids may correspond with the attendance preference of ants (Völkl et al. 1999; Woodring et al. 2004; Xu and Chen 2021). It could be interesting to test whether honeydew quality and composition is associated with the observed ant preference hierarchy in the root aphid community.

The cues that trigger the transport behavior of the studied root aphids are unknown. Cuticular hydrocarbons may differ between ant-tended and untended aphids (Lang and Menzel 2011; Sakata et al. 2017). However, it is not clear whether hydrocarbon blends also trigger carrying behavior in mutualistic aphids. Very specialized parasites of ants are also transported to the nest (Hölldobler 1967; Elmes et al. 1991; Cammaerts 1999; Hölldobler et al. 2018). In these parasites, the mimicking of the cuticular profile of the larva has been invoked as the mechanism to trigger carrying behavior (Elmes et al. 1991). Likewise the transport of a parasitic morph of the root aphid *Paracletus cimiciformis* is likely induced by the resemblance of the cuticular hydrocarbon profile to the host larva (Salazar et al. 2015). The cues that induce seed transport in ants are different. Researchers could demonstrate with bioassays that the fatty acid fraction from the elaiosome stimulates the transport behavior (Marshall et al. 1979; Skidmore and Heithaus 1988; Hughes et al. 1994). Further research should explore whether the transport of root aphids is caused by mimicking the pheromones of the host larva or by producing a unique blend of hydrocarbons or other attractive compounds, as observed in myrmecochorous seeds.

It is puzzling how different root aphids that compete for the services of their host *L. flavus* can co-exist in the same nest environment. As they show different degrees of attractiveness, one could expect that the aphids with the lowest attractivity would be outcompeted if no stabilizing mechanisms would operate (Johnson and Bronstein 2019). Coexistence of mutualists, however, may be promoted through niche partitioning (Sampayo et al. 2007; Palmer et al. 2012). In the case of the subterranean root aphid community, this is favored by the preference for different host plants in the nest, with *Trama* being specialized on *Taraxum* roots, whereas the others on grass roots. In addition, root aphids may prefer spatially different sites of the root network of a single plant, analogous to aboveground aphids that are known to target different sites of a plant (Völkl 1989; Inbar and Wool 1995). During opening of the nests, it appeared that the root aphid *F. formicaria* prefers roots just under the soil level, whereas other aphids could be found much deeper in the nest. Other plausible stabilizing mechanisms are variability in resource acquisition, competition colonization trade-offs and the association with alternative host species. In line with the seed preferences in myrmecochorous ants (Leal et al. 2014), the specialist *L. flavus* may be more selective for its partners than more generalist and co-occurring ants such as *L. niger*. Lastly, the *L. flavus* host may cull the most dominant aphid by feeding on its nymphs (Ivens et al. 2012).

Overall, this study demonstrates that the root aphid-ant mutualism involves disparate transport services which may result in competitive inequalities in the guild of aphids. This multispecies mutualism is a promising system to test different rules of community assembly in symbiont guilds and to explore which factors drive the preference hierarchies of both host and symbiont.

## Appendix. Supplementary data

ESM1: Fig. S1. Map with the sampled nests for the choice and no-choice experiment

ESM2: Video S2. Collection and transport of root aphids in the experimental setup.

ESM3: raw data

