## Supplementary material for "Differential transport of a guild of mutualistic root aphids by the ant *Lasius flavus*": Fig. S1

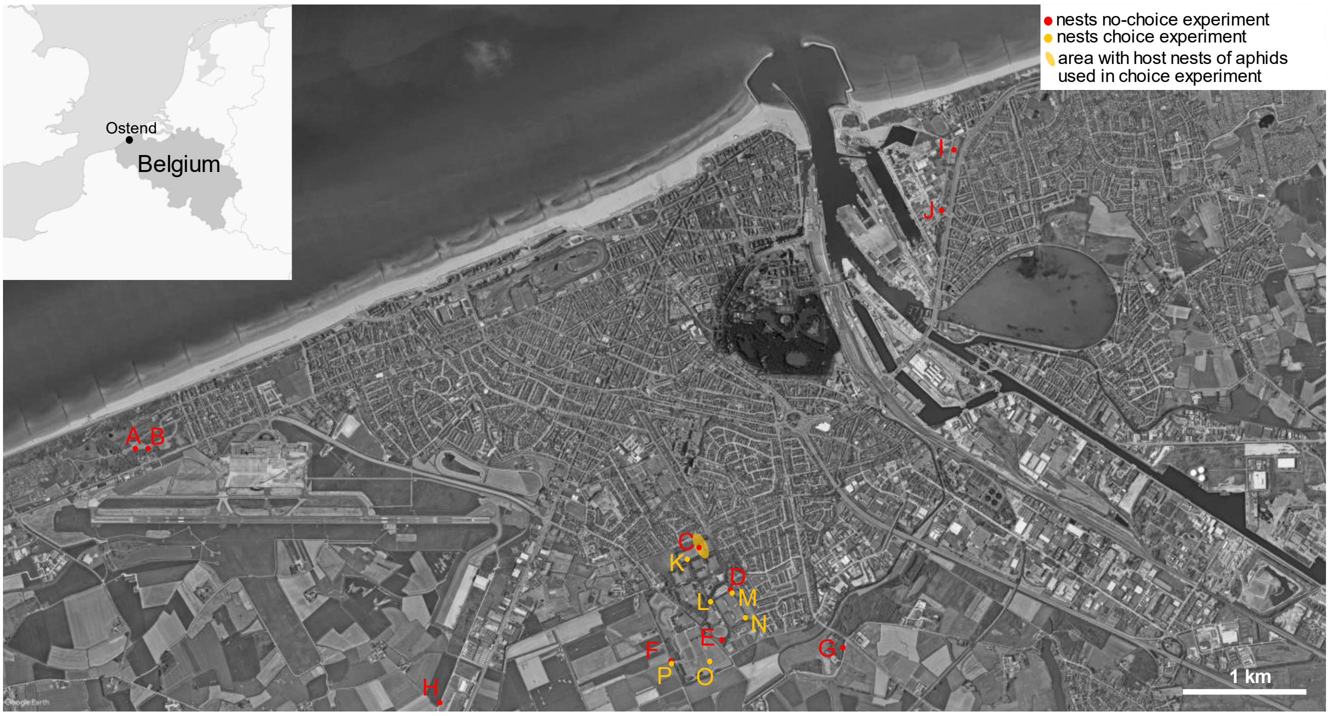


Fig. S1. Overview of the sampled nests in the study area in and around the city of Ostend, Belgium. Nests A – J used in no-choice experiment, nests K – P used in choice experiment.
